# Fast skeletal myosin binding protein-C expression exacerbates dysfunction in heart failure

**DOI:** 10.1101/2024.04.30.591979

**Authors:** James W. McNamara, Taejeong Song, Perwez Alam, Aleksandra Binek, Rohit R. Singh, Michelle L. Nieman, Sheryl E. Koch, Malina J. Ivey, Thomas L. Lynch, Jack Rubinstein, J-P Jin, John N. Lorenz, Jennifer E. Van Eyk, Onur Kanisicak, Sakthivel Sadayappan

## Abstract

During heart failure, gene and protein expression profiles undergo extensive compensatory and pathological remodeling. We previously observed that fast skeletal myosin binding protein-C (fMyBP-C) is upregulated in diseased mouse hearts. While fMyBP-C shares significant homology with its cardiac paralog, cardiac myosin binding protein-C (cMyBP-C), there are key differences that may affect cardiac function. However, it is unknown if the expression of fMyBP-C expression in the heart is a pathological or compensatory response. We aim to elucidate the cardiac consequence of either increased or knockout of fMyBP-C expression. To determine the sufficiency of fMyBP-C to cause cardiac dysfunction, we generated cardiac-specific fMyBP-C over-expression mice. These mice were further crossed into a cMyBP-C null model to assess the effect of fMyBP-C in the heart in the complete absence of cMyBP-C. Finally, fMyBP-C null mice underwent transverse aortic constriction (TAC) to define the requirement of fMyBP-C during heart failure development. We confirmed the upregulation of fMyBP-C in several models of cardiac disease, including the use of lineage tracing. Low levels of fMyBP-C caused mild cardiac remodeling and sarcomere dysfunction. Exclusive expression of fMyBP-C in a heart failure model further exacerbated cardiac pathology. Following 8 weeks of TAC, fMyBP-C null mice demonstrated greater protection against heart failure development. Mechanistically, this may be due to the differential regulation of the myosin super-relaxed state. These findings suggest that the elevated expression of fMyBP-C in diseased hearts is a pathological response. Targeted therapies to prevent upregulation of fMyBP-C may prove beneficial in the treatment of heart failure.

**Significance Statement:** Recently, the sarcomere – the machinery that controls heart and muscle contraction - has emerged as a central target for development of cardiac therapeutics. However, there remains much to understand about how the sarcomere is modified in response to disease. We recently discovered that a protein normally expressed in skeletal muscle, is present in the heart in certain settings of heart disease. How this skeletal muscle protein affects the function of the heart remained unknown. Using genetically engineered mouse models to modulate expression of this skeletal muscle protein, we determined that expression of this skeletal muscle protein in the heart negatively affects cardiac performance. Importantly, deletion of this protein from the heart could improve heart function suggesting a possible therapeutic avenue.

## INTRODUCTION

Heart failure is a devastating global disease with growing prevalence that places extreme burden on patients, families, and communities (1). While recent progress has improved outcomes, treatments remain insufficient for a disease of this scale (2). There is thus an urgent requirement for the development of new and more effective therapies based on the underlying molecular mechanisms of cardiac disease.

During the onset of heart failure, there are dramatic alterations in transcriptional and proteomic networks in the heart (3-6). These alterations include activation of fetal (6, 7), pro-fibrotic (8, 9), immune (10, 11), and pro-hypertrophic (12, 13) programs. While these networks are complex and variable, they ultimately converge on a common endpoint which results in pathologically diminished contractile function (14). Accordingly, recent years have seen an increased focus towards the development of sarcomere-targeted therapies for the treatment of cardiac diseases(15). Clearly an in-depth understanding of the molecular alterations at the sarcomere are key to success of such drug development.

We and others have previously observed the upregulation of the skeletal sarcomeric gene *Mybpc2*, encoding fMyBP-C, in diseased hearts (16-18). The myosin binding protein-C (MyBP-C) family are thick filament accessory proteins which regulate myocyte contractility via transient activation of thin filaments and inhibition of thick filaments (Fig. 1A) (19-24). fMyBP-C is almost exclusively expressed in fast-type muscles (25), while the adult heart expresses almost exclusively cardiac MyBP-C (cMyBP-C) (26). Although these proteins share significant homology, there are key differences which may impart functional differences in the heart, including absence of regulatory phosphorylation sites and divergence of N-terminal domains (26, 27). *In vitro* data from our previous studies has suggested fMyBP-C expression may induce cardiac dysfunction (28), but it is unknown whether its expression is a compensatory or pathological response *in vivo*.

**Figure 1.**
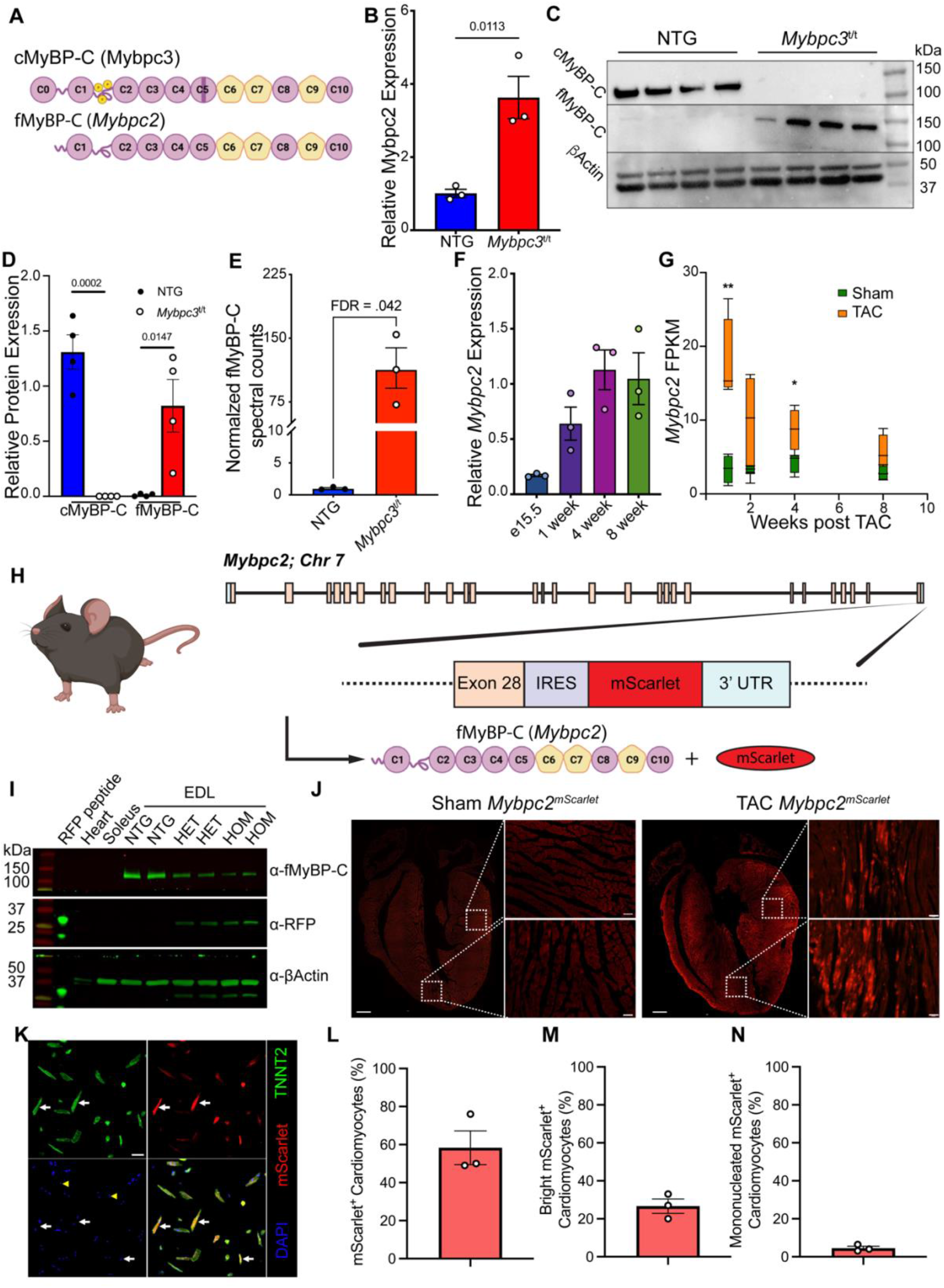
Upregulation of *Mybpc2* in response to cardiac stress. A) Comparison of the domain structures of cMyBP-C and fMyBP-C. Circles represent immunoglobulin-like domains and pentagons fibronectin-like domains. Functionally annotated phosphorylation sites are indicated by “p,” and the cardiac-specific C5 insertion is demonstrated by the darkened rectangle. B) Gene expression analysis of *Mybpc2* in NTG and *Mybpc3*^*t/t*^ mouse hearts (n=3 per group). C) and D) Western blot analysis of cMyBP-C and fMyBP-C in NTG and *Mybpc3*^*t/t*^ mouse hearts (n=4 per group). E) Mass spectrometry spectral counts of fMyBP-C in NTG and *Mybpc3*^*t/t*^ ventricles (n=3 per group). F) *Mybpc2* expression in healthy mouse ventricle across developmental timepoints (n=3 per group). G) Transcriptional analysis of *Mybpc2* expression following TAC (data obtained from Geodata set GSE66630, n=4 per group). H) Gene targeting strategy for *Mybpc2*^mScarlet^ mice. I) Western blot demonstrating specific expression of mScarlet to fast muscle type in naïve *Mybpc2*^mScarlet^ mice. J) Representative immunofluorescent labeling of Sham and TAC *Mybpc2*^mScarlet^ mice. K) Immunofluorescent labeling of enzymatically digested cardiomyocytes from *Mybpc2*^mScarlet^ mice following TAC. L)-N) Quantification of percent of mScarlet positive cardiomyocytes following TAC in *Mybpc2*^mScarlet^ mice (n=3). Data is expressed as mean ± SEM, except in F) in which minimum to maximum values are shown. * Represents p<0.05, ** p<0.01, and *** p<0.001.

In this study, we address the functional consequences of fMyBP-C expression in heart failure, we generated cardiac-specific *Mybpc2* transgenic (*Mybpc2*^Tg^) mice. We observed mild hypertrophy and sarcomeric dysfunction in the absence of global cardiac dysfunction. This model was further crossed into a model of heart failure (*Mybpc3*^*t/t*^) and found that fMyBP-C expression exacerbated the cardiac phenotype. Finally, we performed transverse aortic constriction to induce heart failure in *Mybpc2* knockout (*Mybpc2*^KO^) mice. These mice were exhibited greater resistance to heart failure. Overall, this study demonstrates the upregulation of fMyBP-C in heart failure may be a maladaptive process which further exacerbates cardiac dysfunction.

## RESULTS

### Upregulation of fMyBP-C in cardiac disease

To establish the framework for this study, we first aimed to validate our previous study which demonstrated fMyBP-C expression is increased in models of cardiac disease (16). In 8-week-old *Mybpc3*^t/t^ hearts, quantitative reverse transcriptase-PCR (qRT-PCR) revealed *Mybpc2* expression was increased nearly four-fold compared to NTG controls (Fig. 1B). Western blot analyses further demonstrated the elevated expression of fMyBP-C protein within diseased myocardium of *Mybpc3*^t/t^ mice (Fig.1C, D). Mass spectrometry was used to validate this increase of the fMyBP-C protein quantity in the left ventricles of *Mybpc3*^t/t^ mice, demonstrating more than 100-fold induction (Fig. 1E). Since re-expression of some skeletal muscle isoforms of muscle genes has previously been associated with fetal gene reprogramming in heart disease (29), we performed qRT-PCR on mouse hearts at e15.5, 1-, 4-, and 8-weeks old. Expression of *Mybpc2* mRNA was not elevated in fetal hearts (Fig. 1F), suggesting that this response may be distinct from the fetal gene program. We next investigated the expression of *Mybpc2* mRNA following transverse aortic constriction (TAC) surgery. Interrogation of publicly available RNAseq data (30), demonstrated the increased expression of *Mybpc2* in TAC hearts, which gradually decreased with time (Fig. 1G). To confirm this finding, we performed lineage tracing of *Mybpc2* using reporter mouse (*Mybpc2*^*mScarlet*^) to express the fluorescent protein, mScarlet, under the control of the endogenous *Mybpc2* locus (Fig. 1H, I, Fig. S1). These mice were subjected to TAC surgery followed by histological assessment of reporter activation. Following TAC, *Mybpc2*^*mScarlet*^ hearts showed clear expression of mScarlet in cardiac cells (Fig. 1J). A closer examination from enzymatically digested hearts confirmed the expression of mScarlet within most of the cardiomyocytes of *Mybpc2*^*mScarlet*^ mice following TAC surgery (Fig. 1K-N). Together, these data demonstrate the expression of *Mybpc2* may be aberrantly expressed in ventricular muscle and may associate with cardiac dysfunction in certain settings.

### Cardiac specific expression of fMyBP-C induces subclinical cardiac dysfunction

To determine the consequence of increased fMyBP-C expression in the heart, we generated novel transgenic mice (*Mybpc2*^Tg^) to specifically overexpress *Mybpc2* in cardiomyocytes, driven by the α-myosin heavy chain promoter (Fig. 2A). Western blot analysis demonstrated the expression of fMyBP-C within the myofilament fraction of *Mybpc2*^Tg^ hearts (Fig. 2B), with a small, but significant decrease in cMyBP-C protein quantity (Fig. 2C). Importantly, the decrease in cMyBP-C protein was comparable to the increase in fMyBP-C, maintaining total MyBP-C protein quantity (Fig. 2D, E, Fig. S2). Further, fMyBP-C correctly localized to the A-bands of enzymatically digested *Mybpc2*^Tg^ cardiomyocytes (Fig. 2F-I). qRT-PCR established that while *Mybpc2* mRNA expression was highly increased in *Mybpc2*^Tg^ hearts, there was no change in *Mybpc3 levels* (Fig. 2J, K), suggesting that fMyBP-C replacement of cMyBP-C occurs post-translation. Mass spectrometry was performed on the ventricles of NTG and *Mybpc2*^Tg^ mice to assess changes in the proteome due to increased fMyBP-C protein quantity. Of the 3313 proteins quantified only 5% were modified (143 proteins were significantly reduced, 34 were increased); and the majority of these proteins were not significantly enriched in any gene ontology pathways (Figure 2M). Indeed only fatty acid metabolism was moderately enriched in downregulated proteins (Fold-enrichment 3.04, FDR = 4.51x10^−2^). Furthermore, hierarchical clustering of differentially expressed proteins between NTG, *Mybpc2*^Tg^, and *Mybpc3*^*t/t*^ further supported that the expression of fMyBP-C in otherwise healthy ventricles does not substantially impact the cardiac proteome (Fig. 2N). Furthermore, no major histopathology of *Mybpc2*^Tg^ hearts was observed, except for a mild dilatation of both ventricles (Fig. 2L).

**Figure 2.**
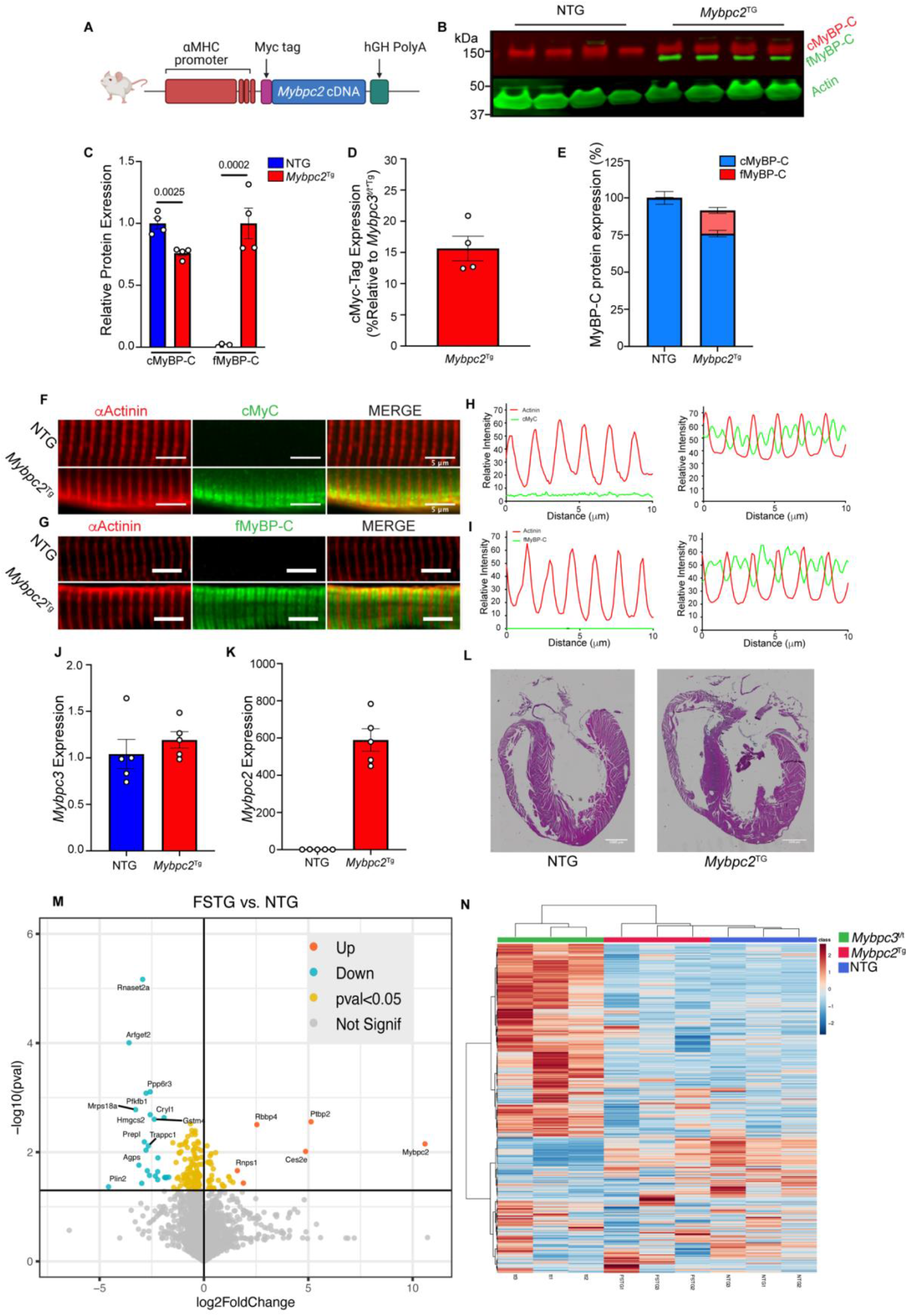
Generation of a mouse model to overexpress fMyBP-C in the heart. A) Strategy for cardiac-specific over-expression of *Mybpc2*. B) and C) Western blot and quantification of cMyBP-C and fMyBP-C expression in NTG and *Mybpc2*^Tg^ mouse hearts (n=4 per group). D) Western blot quantitation of transgene expression via benchmarking cMyc fusion tag expression to mouse hearts expressing 100% cMyc tagged cMyBP-C (n=4). E) Total MyBP-C expression levels determined by Western blot. Immunofluorescent imaging of isolated cardiomyocytes stained for F) alpha-actinin and cMyc-tag and G) alpha-actinin and fMyBP-C. H) and I) Line scans corresponding to confocal images in F) and G), respectively. qRT-PCR analysis of J) *Mybpc3* and K) *Mybpc2* expression in in NTG and *Mybpc2*^Tg^ mouse hearts (n=5 each). L) Representative histological section of in NTG and *Mybpc2*^Tg^ mouse heart. M) Volcano plot showing proteins that are differentially expressed between NTG and *Mybpc2*^Tg^ mouse heart as measured by mass spectrometry. N) Heatmap showing hierarchical clustering of differentially expressed proteins between NTG, *Mybpc2*^Tg^, and *Mybpc2*^t/t^. Data is expressed as mean ± SEM. P-values are displayed within figure.

We next defined the functional consequences of fMyBP-C expression in the heart. Cardiac function of 12-week-old mice were analyzed by echocardiography (Supplementary Table 1) and invasive hemodynamics. Heart weight-to-body weight (Fig. 3A) was significantly increased in *Mybpc2*^Tg^ mice, in the absence of changes in ejection fraction (Fig. 3B). Echocardiography demonstrated increased cardiac dimensions during both systole and diastole (Fig. 3C, D), while *in vivo* pressure measurements were unchanged at baseline and with dobutamine challenge (Fig. 3E). Together, these data demonstrate that fMyBP-C expression in the heart promotes mild ventricular enlargement in the absence of functional changes at the organ level.

**Figure 3.**
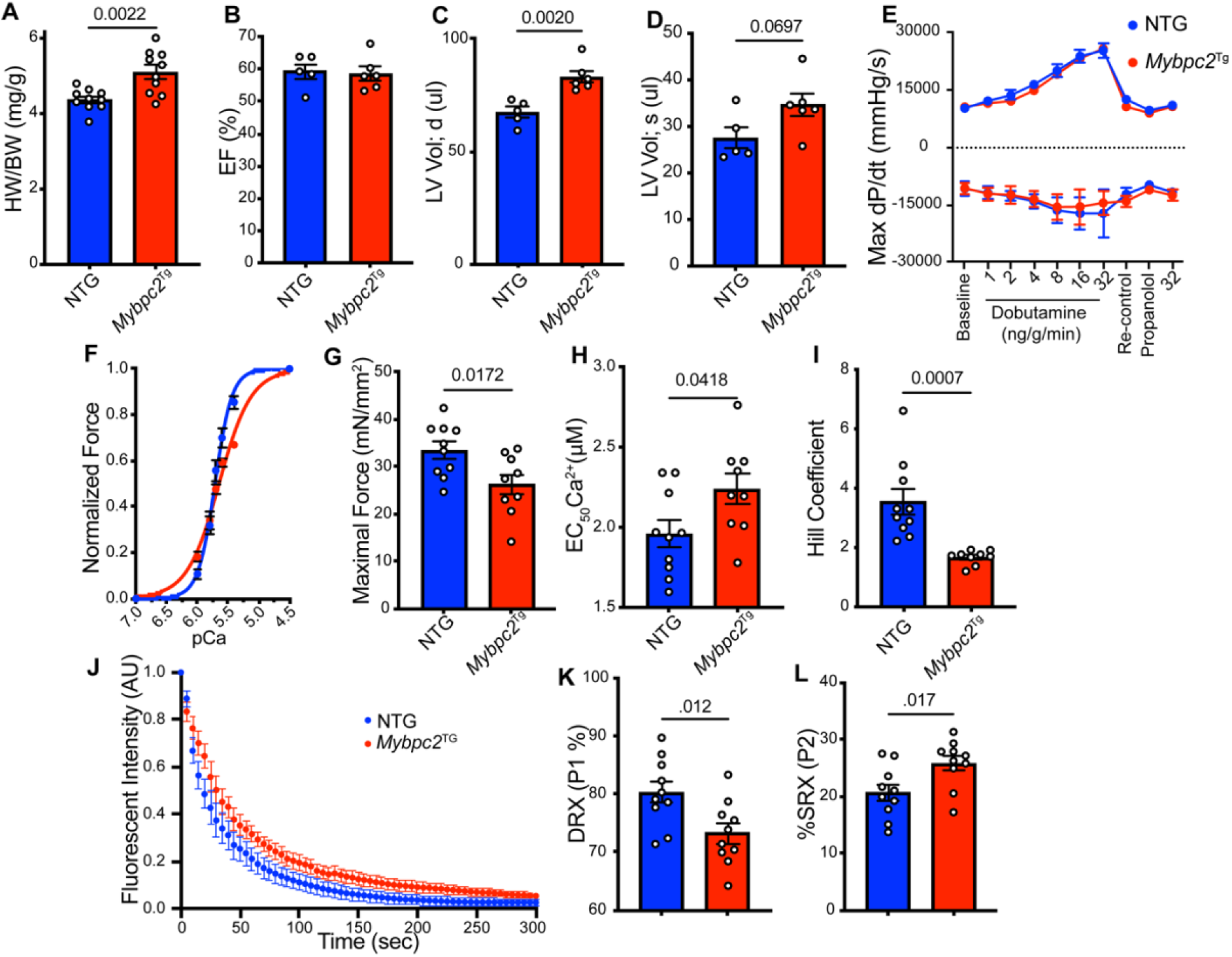
Dysregulation of sarcomere function due to expression of fMyBP-C in the heart. A) 16-week-old heart weight to body weight ratios (n=10 per group). 16-week-old B) LV EF (%); C) diastolic LV volume; and D) systolic LV volume (n=5 for NTG, 6 for *Mybpc2*^Tg^). E) Intraventricular pressure catheterization with dobutamine stimulation (n=4 for NTG, 5 for *Mybpc2*^Tg^). F) Normalized force production of skinned papillary muscles (n=10 fibers for NTG, 9 for. *Mybpc2*^Tg^). G) Maximal force production; H) calcium sensitivity; and I) Hill Coefficient of skinned papillary muscles. J) mantATP nucleotide turnover assay (n=10 fibers each). K) Percentage of DRX and L) SRX myosin heads. Intact cardiomyocyte. Data is expressed as mean ± SEM. P-values are displayed within the figure.

Skinned papillary muscle fibers were used for force-pCa assays to assess subclinical deficits in contractile function (Fig. 3F). *Mybpc2*^Tg^ muscle fibers produced significantly less active force than NTG, concurrent with the desensitization of myofilaments to calcium (Fig. 3G, H). Notably, *Mybpc2*^Tg^ muscle fibers exhibited a lower Hill constant (Fig. 3I), suggesting the co-operativity of muscle activation is blunted upon expression of fMyBP-C. We also observed that skinned *Mybpc2*^Tg^ muscles exhibit greater stability of myosin in the super-relaxed (SRX) state (Fig. 3J-L). It is unclear if this is due to myosin binding differences between fMyBP-C and cMyBP-C or a total reduction in MyBP-C phosphorylation, known to stabilize the SRX (21, 31). Collectively, these data suggest that low level expression of fMyBP-C in the heart is sufficient to drive subclinical deficits in myofilament contractility. Mechanistically, this likely arises from greater stability of myosin heads in the SRX, which subsequently blunt muscle activation *in situ*.

### Complete fMyBP-C replacement exacerbates ventricular dysfunction

Having demonstrated that incorporation of fMyBP-C into the cardiac sarcomere causes mild dysfunction, we investigated the consequence of 100% expression of fMyBP-C in the mouse heart. To achieve this, we crossed the *Mybpc2*^Tg^ with a cMyBP-C null mouse model (*Mybpc3*^t/t^). Western blots demonstrated that fMyBP-C expression in double transgenic (dTG) mice was equivalent to cMyBP-C in healthy mice (Fig. 4A, B). Echocardiography was performed on 12-week-old mice (Supplementary Table 2), which demonstrated that fMyBP-C overexpression was sufficient to exacerbate cardiac dysfunction in this established model (Fig. 4C). Similar to *Mybpc2*^*Tg*^ mice, the left ventricular chamber dimension in dTG mice was enlarged (Fig. 4D, E), while interventricular septal thickness was significantly reduced (Fig. 4F, G). We further observed greater fibrosis (Fig. 4H, I) in the dTG model compared to *Mybpc3*^t/t^ mice, as well as increased cardiomyocyte hypertrophy (Fig. 4H, J). While *Mybpc2* expression was still 4-fold elevated in *Mybpc3*^t/t^ mice compared to NTG, this was more prominent in dTG mice (Fig. 4K), while *Mybpc3* expression was reduced in both these mice. Furthermore, we also observed elevated levels on *Nppa* only in dTG mice, while *Nppb* was higher in both *Mybpc3*^t/t^ and dTG mice compared to controls. Counterintuitively, this data suggested that the expression of fMyBP-C in the heart is worse than the complete lack of MyBP-C expression.

**Figure 4.**
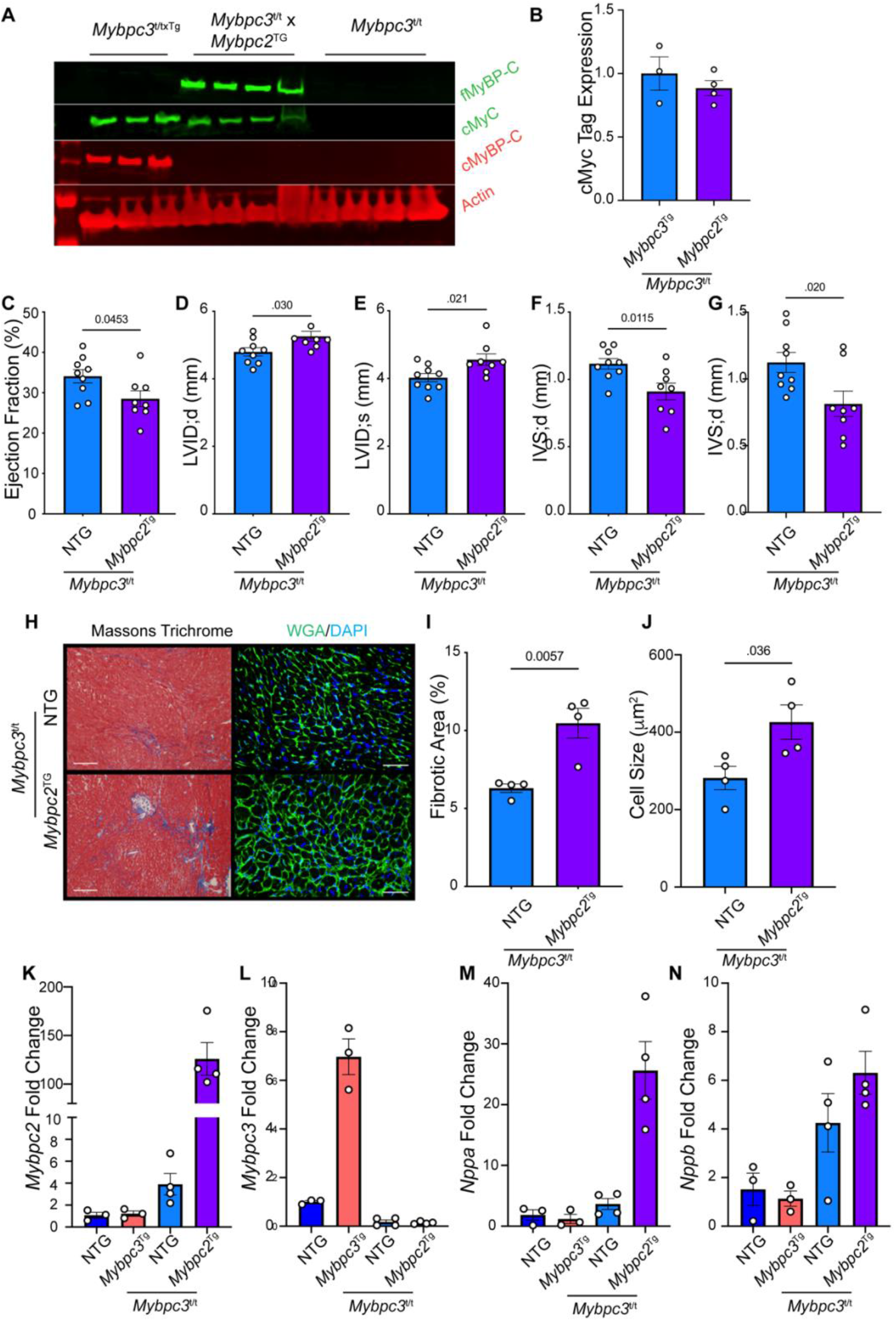
Complete replacement of cMyBP-C with fMyBP-C exacerbates cardiac pathology. A) Western blot comparing fMyBP-C overexpression in dTG to control. B) Quantification of cMyc tag expression between *Mybpc3*^*t/t x Tg*^ (n=3) and dTG (n=4). C)-G) Echocardiographic parameters (n=9 for *Mybpc3*^*t/t*^; 8 for dTG). H) Masson’s trichrome and wheat germ agglutinin. I) Fibrosis and J) cell size quantification (n=4 per group). K)-N) qRT-PCR expression of *Mybpc2, Mybpc3, Nppa*, and *Nppb (*n=3-4 per group). Data is expressed as mean ± SEM. P-values are displayed within figure.

The exacerbation of heart function upon 100% expression of fMyBP-C was further investigated using skinned ventricular preparations and isolated ventricular myocytes. Skinned fibers were subjected to force-pCa (Fig. 5A) and contractile kinetics by means of a slack re-stretch protocol. Maximal force production (Fig. 5B) was significantly reduced in dTG mice compared to control mice (*Mybpc3* overexpressed in the *Mybpc3*^*t/t*^ background). No differences were observed in calcium sensitivity (Fig.5C) or Hill co-efficient (Fig. 5D) between groups. To assess the rate of contraction, slack re-stretch assays were performed at an intermediate and maximal calcium concentration. In line with established results(32), *Mybpc*^*t/t*^ mice exhibited accelerated tension redevelopment (*k*_TR_) at submaximal calcium activation (Fig. 5E) which normalized at maximal calcium levels (Fig. 5F). Conversely, the submaximal *k*_TR_ of dTG preparations was comparable to control mice (Figure 5E) but could not normalize at maximal levels (Fig. 5F). This data collectively highlights key functional differences in the cardiac and fast skeletal paralogs of MyBP-C which may differentially impact cellular contractility and emphasizes the inability of fMyBP-C to functionally replace its cardiac paralog.

**Figure 5.**
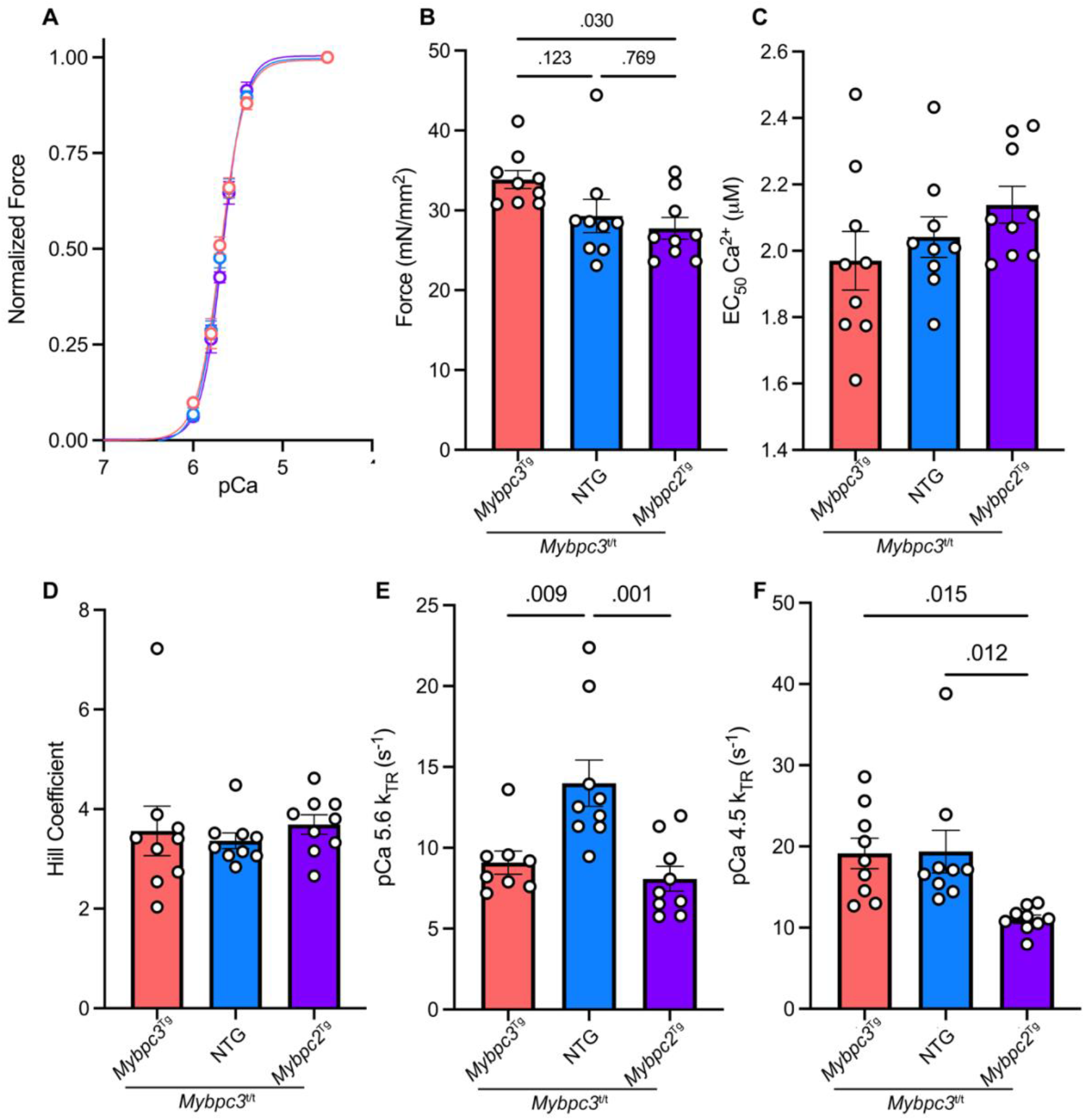
Impaired force and kinetics in cardiac muscle expressing only fMyBP-C. A) Normalized force production of skinned papillary muscles. B) Maximal force production; C) calcium sensitivity; and D) Hill Coefficient from skinned papillary muscles. k_TR_ measured at E) intermediate and F) maximal calcium concentration (n=8-9 fibers per group). Enzymatically isolated cardiomyocyte contractility with or without OM: G) cell shortening; H) contraction and I) relaxation velocities. The dotted line indicates values for NTG (-) OM. Data is expressed as mean ± SEM. P-values are displayed within the figure.

### Deletion of *Mybpc2* diminishes pressure overload-induced heart failure

Having demonstrated that fMyBP-C expression may be detrimentally induced in specific forms of cardiac stress, we questioned whether its absence could ameliorate cardiac remodeling following pressure overload. Eight-week-old control or *Myb*pc2^KO^ mice, which have a normal cardiac function (25), were subjected to TAC for 8 weeks followed by echocardiography. Normalized heart weights were significantly elevated following TAC in both WT and *Myb*pc2^KO^ mice compared to sham mice (Fig. 6A). However, this effect was less prominent in *Myb*pc2^KO^ than WT. Similarly, reductions to ejection fraction and fractional shortening were significantly diminished following TAC in *Myb*pc2^KO^ mice compared to WT mice (Fig. 6B, C). Genetic deletion of *Mybpc2* similarly resulted amelioration of detrimental reductions in stroke volume, cardiac output, and wall thickening (Fig. 6D-F). Strikingly, we observed a significant stabilization of the myosin SRX state in NTG mice following TAC surgery, which was partially ameliorated in absence on *Mybpc2* expression (Fig. 6G-I).

**Figure 6.**
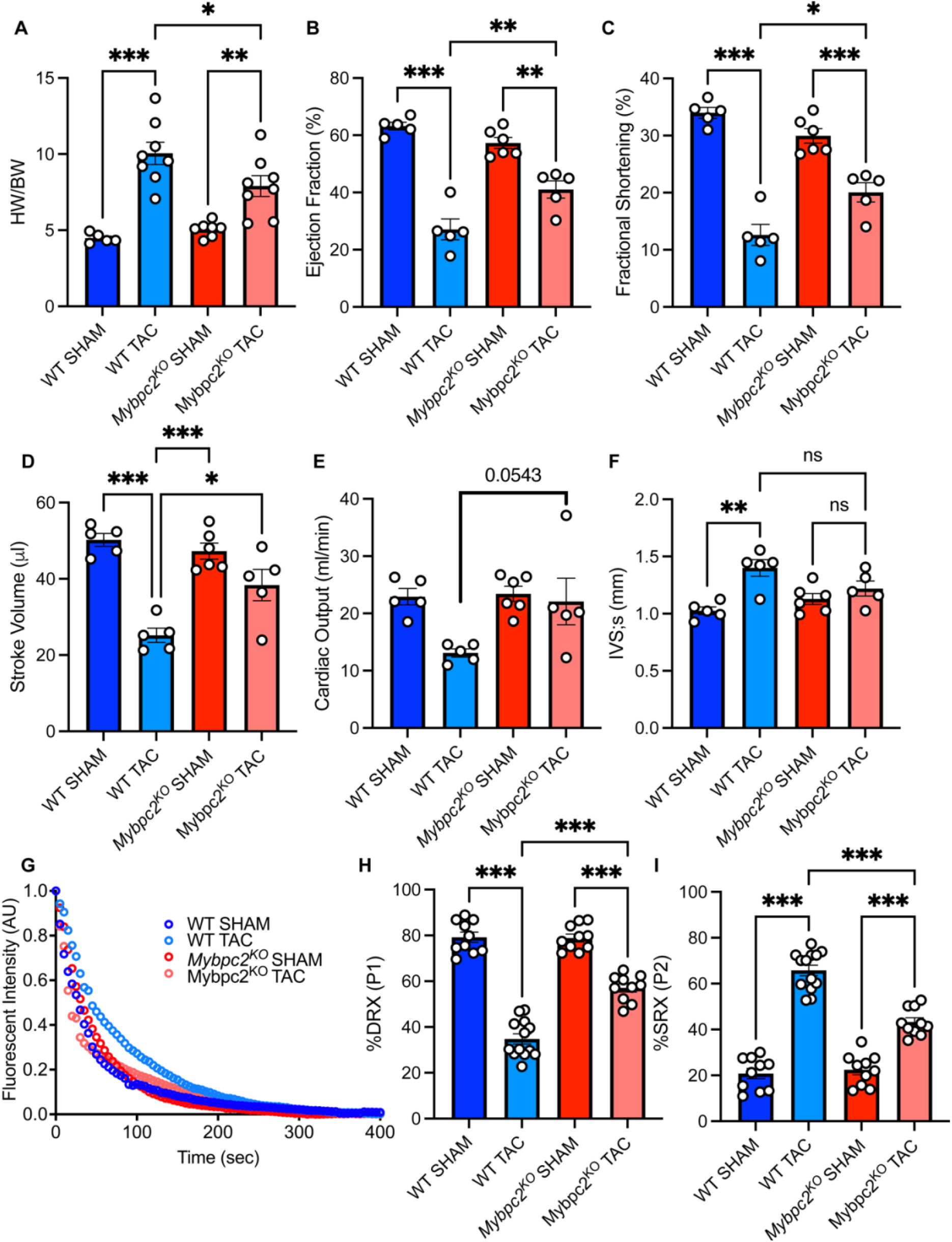
Deletion of *Mybpc2* ameliorates dysfunction in response to pressure overload. A) Heart weight to body weight ratios. B-F) Echocardiographic parameters of WT and *Mybpc2*^-/-^ sham and TAC mice. G) mantATP nucleotide turnover assay experiments from skinned ventricular muscle from WT and *Mybpc2*^-/-^ sham and TAC mice. Quantification of H) DRX and I) SRX populations of myosin from Single ATP turnover experiments. Data is expressed as mean ± SEM * Represents p<0.05, ** p<0.01, and *** p<0.001.

## DISCUSSION

In this study, we used multiple genetic mouse models to assess the functional effect of elevated myocardial expression of *Mybpc2* during cardiac stress. Three *Mybpc* genes are expressed in the mammalian genome-slow, fast, and cardiac. These encode proteins with substantial structural homology. consisting largely of immunoglobulin- and fibronectin-like domains (Fig. 1A) (28). It has long been understood that the cardiac and skeletal paralogs do not exhibit isoform transcomplentation in healthy hearts (33, 34). However, our group has recently demonstrated that ectopic expression of *Mybpc2*, the gene encoding the fast paralog of MyBP-C is upregulated in the heart of mice lacking *Mybpc3 (16)*. This finding has also been observed in other models of cardiac stress, including dilated and hypertrophic cardiomyopathy, and pressure overload (17, 18, 30). The mechanisms regulating the increased expression of *Mybpc2* in the hearts of diseased mice remain unclear. However, it may not be surprising that there may be paralog shifts in certain settings. Indeed, many sarcomeric proteins, exhibit shifts in skeletal and cardiac protein expression in the heart, discussed below. However, it is unclear whether, and if so, how, the expression of fMyBP-C in the ventricular muscle affects cardiac function.

We first validated the expression of fMyBP-C in cMyBP-C null mice. Importantly, unbiased mass spectrometry confirmed a roughly hundred-fold induction of fMyBP-C expression in the ventricles of cMyBP-C null mice compared to non-transgenic mice. Using genetic lineage tracing of *Mybpc2* we found that pressure overload of the left ventricle also induce expression of *Mybpc2*. These provide clear evidence that fMyBP-C is upregulated in response to multiple forms of cardiac stress. We next demonstrated in cardiomyocyte-specific transgenic mice that low-level expression of fMyBP-C (the protein product of *Mybpc2*) caused a mild increase in cardiac dimensions, in the absence of overt disease. This was sufficient, however, to cause molecular defects in contractile function as measured in skinned papillary muscles. Somewhat surprisingly, the complete replacement of cMyBP-C with fMyBP-C was more detrimental to heart function than the absence of any MyBP-C molecule. Finally, deletion of *Mybpc2* in mice could partially ameliorate cardiac deficits following pressure overload. Mechanistically, we found that the expression, or absence, of fMyBP-C affected the stability of cardiac myosin SRX.

Why *Mybpc2* expression has observed to increase in specific settings of heart disease was unclear. One possibility was that fMyBP-C contributed to the fetal gene program, which results in the re-expression of numerous genes which are expressed during cardiac development (35). In mice, this includes changes to the expression profile of sarcomeric genes. For example, *Myh7* is the primary sarcomeric motor protein expressed in the fetal, but not adult, mouse heart. After cardiac insult *Myh7* expression is reactivated (36). Similarly, the N2B:N2BA titin favors the expression of the compliant N2BA isoform which is expressed at higher levels during development. Further, several skeletal muscle genes have been implicated in the fetal gene response including skeletal isoforms of actin and troponin-I (29). It is thought that these modifications are compensatory responses to metabolic reprogramming of the diseased heart. Indeed, beta myosin heavy chain hydrolyses ATP less rapidly than the alpha isoform. Thus, the expression of this slower isoform may act to increase metabolic efficiency of the diseased heart, while imparting marginal reductions in contractile performance (37). Similarly, slow skeletal troponin I confers greater protection to the heart during stressful periods, such as acidosis, while also reducing sarcomeric ATPase (38-40). It was possible that the expression of fMyBP-C may be related to these compensatory fetal gene mechanisms in attempt to conserve cardiac performance.

However, we and others have not found evidence that fMyBP-C is expressed in the fetal heart. These data suggest an alternative mechanism behind the increased fMyBP-C expression in disease. We propose this may be related to the mechanics of the different myosin binding protein-C paralogs. In skeletal muscle, fMyBP-C has specialized expression towards fast, glycolytic myofibers (25). These fibers are activated during high load activities and produce greater force than the oxidative myofibers. It is thought that fMyBP-C contributes to this activation by slowing actin filament sliding though the sarcomere, which in turn co-operatively recruits more force producing actomyosin complexes (41). It is thus possible that fMyBP-C is an initial compensatory mechanism within the myocardium to respond to pathological changes in mechanical stress.

These paralogs do differ, however, in several aspects. For one, the skeletal isoforms of MyBP-C lack the N-terminal “C0” domain that is present in the cardiac protein. This C0 domain can bind the myosin regulatory light chain, and activate the thin filament, providing dual modulation of cardiac muscle activation (42, 43). cMyBP-C has three functionally annotated phosphoserines which are regulated by adrenergic stimuli. These sites are absent in fMyBP-C. Indeed, not no phosphorylation sites have been functionally annotated for fMyBP-C to date. Phosphorylation of cMyBP-C is critical for maintaining normal systolic and diastolic function of the heart (44). These factors likely contribute to the alterations in molecular contractility observed in this study. We observed that the presence of fMyBP-C slows tension development following slack tests at both intermediate and high calcium concentrations, reduces force of contraction, and cellular shortening. cMyBP-C carries intrinsic ability both activate and inhibit actomyosin contractility in a calcium-dependent manner, which fMyBP-C, in contrast, but can inhibit activity at high concentrations (28, 41). Further, adenoviral overexpression of fMyBP-C appeared to affect cellular contractility in enzymatically isolated adult rat cardiomyocytes(28). Our evidence suggest that one possible mechanism may be that fMyBP-C and cMyBP-C differentially regulate the cardiac SRX state of myosin. While it is important to note that the biochemical SRX does not fully translate to the structural “OFF” state of myosin, it is possible that the greater stability of SRX myosin is likely to impair force of contraction and affect the speed of contraction. The structural and regulatory differences listed above between N-termini may be sufficient to promote the observed stabilization of SRX myosin following expression of fMyBP-C. Changes in thin filament activation, or a combination, would also be valid hypotheses for the observed deficits of these mice. X-Ray diffraction analyses of intact or skinned papillary muscles would likely provide greater insight into the molecular mechanism but are outside the scope of this study.

In conclusion, we have demonstrated the pathological role fMyBP-C plays when expressed during cardiac disease. Although *Mybpc2* expression is low in settings of disease, it is well-known that small perturbations to the sarcomeric lattice may have great affects to cardiac performance adding to. We have shown that reducing expression of *Mybpc2* in pathological stress may partially ameliorate dysfunction. This suggests it may be of translational interest with additional study.

## MATERIALS AND METHODS

Detailed methods are provided in the Online Data Supplement.

### Mouse Models

To generate transgenic mice with cardiac-specific expression of fMyBP-C, the fMyBP-C cDNA, *Mybpc2*, was amplified by polymerase chain reaction for cloning downstream of the alpha myosin heavy chain promoter. Briefly, an N-terminal cMyc tag for transgene recognition was cloned in-frame with *Mybpc2* and SalI restriction enzyme sites were included both 5’ and 3’ of the sequence for subcloning into the a-myosin heavy chain promoter (a gift from Dr. Jeffrey Robbins). Clones were screened for correct orientation of insertion and Sanger sequenced to confirm *Mybpc2* sequence. The transgene was liberated from the vector backbone by digestion with NotI for zygote injections. Multiple *Mybpc2* transgenic (*Mybpc2*^Tg^) founders were identified, but only one showed appreciable expression of fMyBP-C in the heart. Global knockout models for *Mybpc2* (25) and *Mybpc3* (45, 46) have been previously described.

## Abbreviations

Mybpc2: fast myosin binding protein-c gene
Mybpc3: cardiac myosin binding protein-c gene
cMyBP-C: cardiac myosin binding protein-c
fMyBP-C: fast myosin binding protein-c
*Mybpc2*^Tg^: α-myosin heavy chain driven fast myosin binding protein-C transgene
*Mybpc3*^*t/t*^: cardiac myosin binding protein-c knockout
*Mybpc2*^KO^: fast myosin binding protein-c knockout
NTG: non-transgenic (wild-type)
TAC: Transverse aortic constriction
SRX: Super-Relaxed state of myosin

## Acknowledgments

Dr. McNamara (Postdoctoral Fellowship, 17POST33630095), Dr. Song (Postdoctoral Fellowship, 19POST34380448 and Career Development Grant, 23CDA1046498), Dr. Lynch (Predoctoral Fellowship, 15PRE22430028), Dr. Alam (AHA-Postdoctoral Fellowship, 20POST35200267 and AHA-Career Development Grant, 23CDA1052132), and Dr. Binek were supported by the American Heart Association (Postdoctoral Fellowship AHA Award Number: 829444). were supported by American Heart Association. Dr. McNamara has been supported by Heart Foundation Future Leader Fellowship (107192), the Medical Research Future Fund (GNT 2024440), and the Murdoch Children’s Research Institute (MCRI) Early Career Researcher Award. Dr. Singh was supported by the Amgen postdoctoral fellowship. Dr. Sadayappan has received support from National Institutes of Health grants (R01 AR079435, R01 AR079477, R01 AR078001, R01 HL130356, R01 HL105826, R38 HL155775 and R01 HL143490), the American Heart Association, Institutional Undergraduate Student (23IAUST1020498), Transformation (19TPA34830084 and 945748) awards, the PLN Foundation (PLN crazy idea) awards and the Cedars-Sinai Proteomics and Metabolomics Core for access to mass spectrometers. The authors thank Christine E. Seidman and Jonathan G. Seidman, Harvard Medical School, Boston, MA 02115, USA, for providing the cMyBP-C^(t/t)^ mouse model.

## Author contributions

J.W.M., T.S., P.A., A.B., R.R.S., M.L.N., M.J.I., T.L.L., J.N.L., J.E.V., O.K. and S.S. designed research; J.W.M., T.S., P.A., A.B., R.R.S., M.L.N., M.J.I., T.L.L. and O.K., performed research; J-P.J., contributed new reagents/analytic tools; J.W.M., T.S., P.A., A.B., R.R.S., M.L.N., M.J.I., T.L.L., J.N.L., J.E.V., O.K. and S.S. analyzed data; and J.W.M., T.S., O.K. and S.S. wrote the paper.

## Data Availability

The mass spectrometry proteomics data have been deposited to the ProteomeXchange Consortium via the PRIDE partner repository with the dataset identifier PXD047728. Data will be made public upon manuscript publication in peer reviewed journal.

Appendix A. Supplementary data

Appendix B. Detailed Materials and Method Sections

## References

1. G. Savarese, L. H. Lund, Global Public Health Burden of Heart Failure. Card Fail Rev 3, 7–11 (2017).

2. G. A. Figtree et al., A call to action for new global approaches to cardiovascular disease drug solutions. Eur Heart J 42, 1464–1475 (2021).

3. D. Reichart et al., Pathogenic variants damage cell composition and single cell transcription in cardiomyopathies. Science 377, eabo1984 (2022).

4. M. J. Previs et al., Defects in the Proteome and Metabolome in Human Hypertrophic Cardiomyopathy. Circ Heart Fail 15, e009521 (2022).

5. M. Li et al., Core functional nodes and sex-specific pathways in human ischaemic and dilated cardiomyopathy. Nat Commun 11, 2843 (2020).

6. N. R. Mehdiabadi et al., Defining the Fetal Gene Program at Single-Cell Resolution in Pediatric Dilated Cardiomyopathy. Circulation 146, 1105–1108 (2022).

7. M. Rajabi, C. Kassiotis, P. Razeghi, H. Taegtmeyer, Return to the fetal gene program protects the stressed heart: a strong hypothesis. Heart Fail Rev 12, 331–343 (2007).

8. J. G. Travers, F. A. Kamal, J. Robbins, K. E. Yutzey, B. C. Blaxall, Cardiac Fibrosis: The Fibroblast Awakens. Circ Res 118, 1021–1040 (2016).

9. X. Fu et al., Specialized fibroblast differentiated states underlie scar formation in the infarcted mouse heart. J Clin Invest 128, 2127–2143 (2018).

10. F. K. Swirski, M. Nahrendorf, Leukocyte Behavior in Atherosclerosis, Myocardial Infarction, and Heart Failure. Science 339, 161–166 (2013).

11. E. Martini et al., Single-Cell Sequencing of Mouse Heart Immune Infiltrate in Pressure Overload–Driven Heart Failure Reveals Extent of Immune Activation. Circulation 140, 2089–2107 (2019).

12. Y. K. Tham, B. C. Bernardo, J. Y. Y. Ooi, K. L. Weeks, J. R. McMullen, Pathophysiology of cardiac hypertrophy and heart failure: signaling pathways and novel therapeutic targets. Arch Toxicol 89, 1401–1438 (2015).

13. M. Hoshijima, K. R. Chien, Mixed signals in heart failure: cancer rules. JClin Invest 109, 849–855 (2002).

14. N. Hamdani et al., Sarcomeric dysfunction in heart failure. Cardiovasc Res 77, 649–658 (2007).

15. C. Morelli, G. Ingrasciotta, D. Jacoby, A. Masri, I. Olivotto, Sarcomere protein modulation: The new frontier in cardiovascular medicine and beyond. Eur J Intern Med 102, 1–7 (2022).

16. B. Lin et al., Cardiac myosin binding protein-C plays no regulatory role in skeletal muscle structure and function. PLoS One 8, e69671 (2013).

17. E. Farrell et al., Transcriptome Analysis of Cardiac Hypertrophic Growth in MYBPC3-Null Mice Suggests Early Responders in Hypertrophic Remodeling. Front Physiol 9 (2018).

18. S. Vakrou et al., Allele-specific differences in transcriptome, miRNome, and mitochondrial function in two hypertrophic cardiomyopathy mouse models. JCI Insight 3 (2018).

19. S. Rahmanseresht et al., The N terminus of myosin-binding protein C extends toward actin filaments in intact cardiac muscle. J Gen Physiol 153 (2021).

20. M. J. Previs, S. Beck Previs, J. Gulick, J. Robbins, D. M. Warshaw, Molecular mechanics of cardiac myosin-binding protein C in native thick filaments. Science 337, 1215–1218 (2012).

21. J. W. McNamara, R. R. Singh, S. Sadayappan, Cardiac myosin binding protein-C phosphorylation regulates the super-relaxed state of myosin. Proc Natl Acad Sci U S A 116, 11731–11736 (2019).

22. T. L. t. Lynch et al., Amino terminus of cardiac myosin binding protein-C regulates cardiac contractility. J Mol Cell Cardiol 156, 33–44 (2021).

23. S. Ponnam, I. Sevrieva, Y. B. Sun, M. Irving, T. Kampourakis, Site-specific phosphorylation of myosin binding protein-C coordinates thin and thick filament activation in cardiac muscle. Proc Natl Acad Sci U S A 116, 15485–15494 (2019).

24. T. Kampourakis, Z. Yan, M. Gautel, Y. B. Sun, M. Irving, Myosin binding protein-C activates thin filaments and inhibits thick filaments in heart muscle cells. Proc Natl Acad Sci U S A 111, 18763–18768 (2014).

25. T. Song et al., Fast skeletal myosin-binding protein-C regulates fast skeletal muscle contraction. Proc Natl Acad Sci U S A 118 (2021).

26. J. W. McNamara, S. Sadayappan, Skeletal myosin binding protein-C: An increasingly important regulator of striated muscle physiology. Arch Biochem Biophys 660, 121–128 (2018).

27. D. Barefield, S. Sadayappan, Phosphorylation and function of cardiac myosin binding protein-C in health and disease. J Mol Cell Cardiol 48, 866–875 (2010).

28. B. L. Lin et al., Skeletal myosin binding protein-C isoforms regulate thin filament activity in a Ca(2+)-dependent manner. Sci Rep 8, 2604 (2018).

29. A. van der Pol, M. F. Hoes, R. A. de Boer, P. van der Meer, Cardiac foetal reprogramming: a tool to exploit novel treatment targets for the failing heart. J Intern Med 288, 491–506 (2020).

30. X. Ma et al., Revealing Pathway Dynamics in Heart Diseases by Analyzing Multiple Differential Networks. PLoS Comput Biol 11, e1004332 (2015).

31. R. W. Kensler, R. Craig, R. L. Moss, Phosphorylation of cardiac myosin binding protein C releases myosin heads from the surface of cardiac thick filaments. Proc Natl Acad Sci U S A 114, E1355–e1364 (2017).

32. F. S. Korte, K. S. McDonald, S. P. Harris, R. L. Moss, Loaded shortening, power output, and rate of force redevelopment are increased with knockout of cardiac myosin binding protein-C. Circ Res 93, 752–758 (2003).

33. F. Fougerousse et al., Cardiac myosin binding protein C gene is specifically expressed in heart during murine and human development. Circ Res 82, 130–133 (1998).

34. M. Gautel, D. O. Fürst, A. Cocco, S. Schiaffino, Isoform transitions of the myosin binding protein C family in developing human and mouse muscles: lack of isoform transcomplementation in cardiac muscle. Circ Res 82, 124–129 (1998).

35. H. Taegtmeyer, S. Sen, D. Vela, Return to the fetal gene program: a suggested metabolic link to gene expression in the heart. Ann N Y Acad Sci 1188, 191–198 (2010).

36. P. Lu et al., Cardiac Myosin Heavy Chain Reporter Mice to Study Heart Development and Disease. Circ Res 131, 364–366 (2022).

37. J. C. Tardiff et al., Expression of the beta (slow)-isoform of MHC in the adult mouse heart causes dominant-negative functional effects. Am J Physiol Heart Circ Physiol 278, H412-419 (2000).

38. A. N. Carley, D. M. Taglieri, J. Bi, R. J. Solaro, E. D. Lewandowski, Metabolic efficiency promotes protection from pressure overload in hearts expressing slow skeletal troponin I. Circ Heart Fail 8, 119–127 (2015).

39. M. L. Alves et al., Early sensitization of myofilaments to Ca2+ prevents genetically linked dilated cardiomyopathy in mice. Cardiovasc Res 113, 915–925 (2017).

40. B. M. Wolska et al., Expression of slow skeletal troponin I in adult transgenic mouse heart muscle reduces the force decline observed during acidic conditions. J Physiol 536, 863–870 (2001).

41. A. Li et al., Skeletal MyBP-C isoforms tune the molecular contractility of divergent skeletal muscle systems. Proc Natl Acad Sci U S A 116, 21882–21892 (2019).

42. C. Risi et al., N-Terminal Domains of Cardiac Myosin Binding Protein C Cooperatively Activate the Thin Filament. Structure 26, 1604-1611.e1604 (2018).

43. J. Ratti, E. Rostkova, M. Gautel, M. Pfuhl, Structure and interactions of myosinbinding protein C domain C0: cardiac-specific regulation of myosin at its neck? J Biol Chem 286, 12650–12658 (2011).

44. P. C. Rosas et al., Phosphorylation of cardiac Myosin-binding protein-C is a critical mediator of diastolic function. Circ Heart Fail 8, 582–594 (2015).

45. B. K. McConnell et al., Dilated cardiomyopathy in homozygous myosin-binding protein-C mutant mice. J Clin Invest 104, 1235–1244 (1999).

46. D. Y. Barefield et al., Ablation of the calpain-targeted site in cardiac myosin binding protein-C is cardioprotective during ischemia-reperfusion injury. J Mol Cell Cardiol 129, 236–246 (2019).

