## Supplementary material for "Fast skeletal myosin binding protein-C expression exacerbates dysfunction in heart failure": Online Supplemental Data

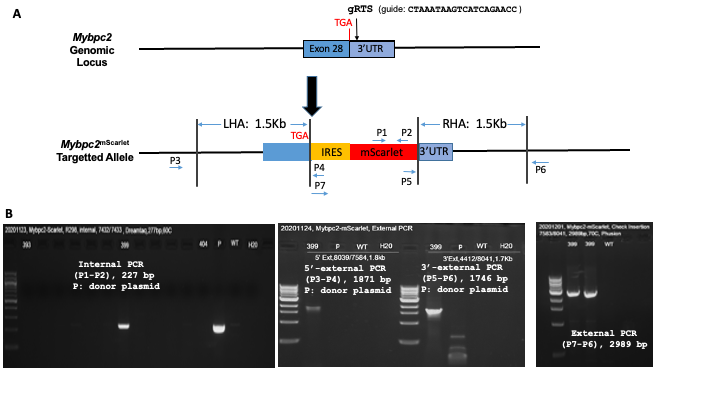

**Supplementary Figure 1. Generation of *Mybpc2*^mScarlet^ mouse model.** A. *Mybpc2*^mScarlet^ targeting strategy. B. Screening of founder mice for *Mybpc2*^mScarlet^ targeting.

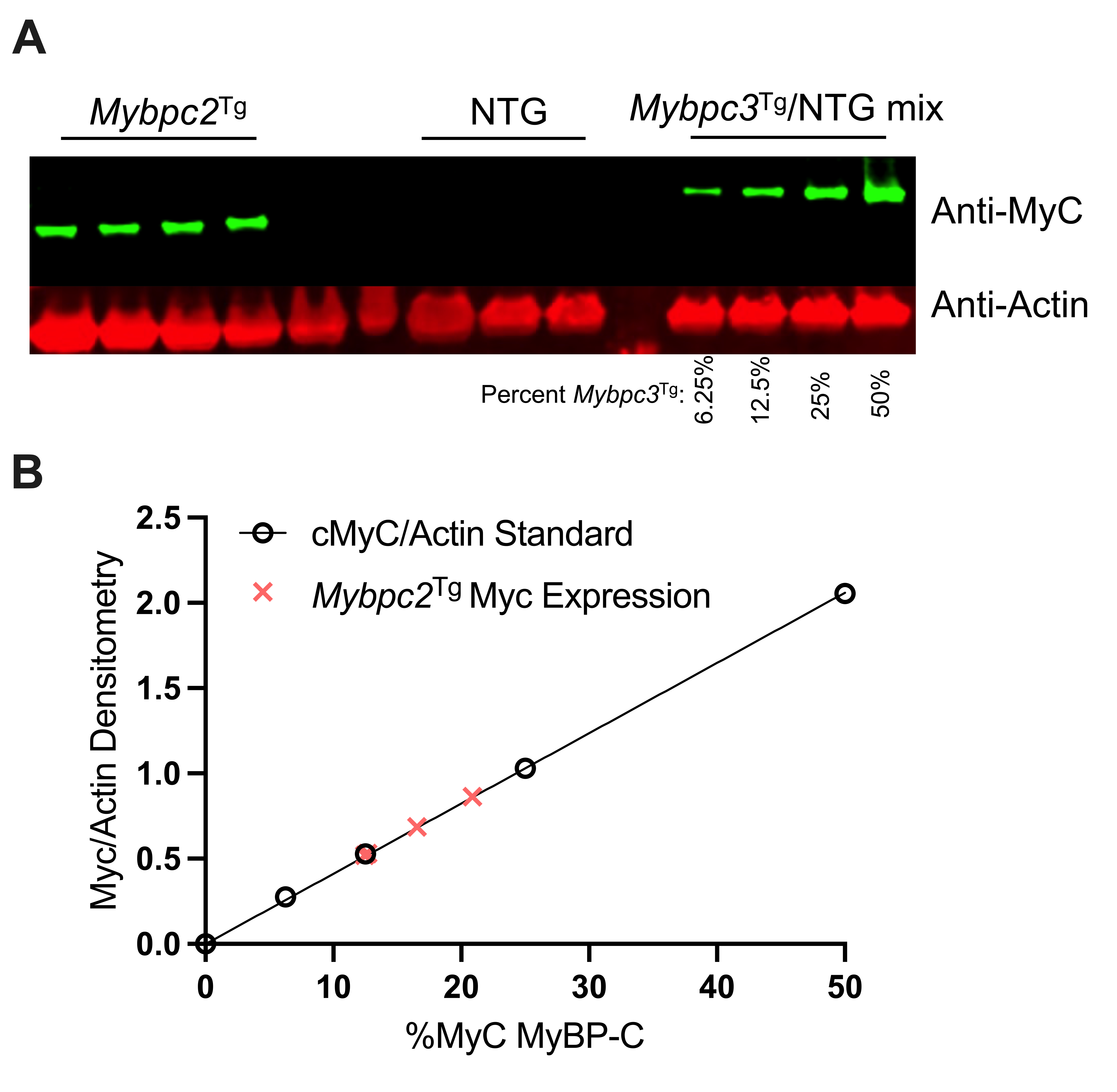

**Supplementary Figure 2. Assessment of relative fMyBP-C amount.** A. Western blot of *Mybpc2*^Tg^ myofilaments compared to a mixture of Myc tagged *Mybpc3*^Tg^ and NTG to assess total fMyBP-C levels. B. Myc-tagged fMyBP-C expression calculated from standard curve of Myc tagged *Mybpc3*^Tg^.

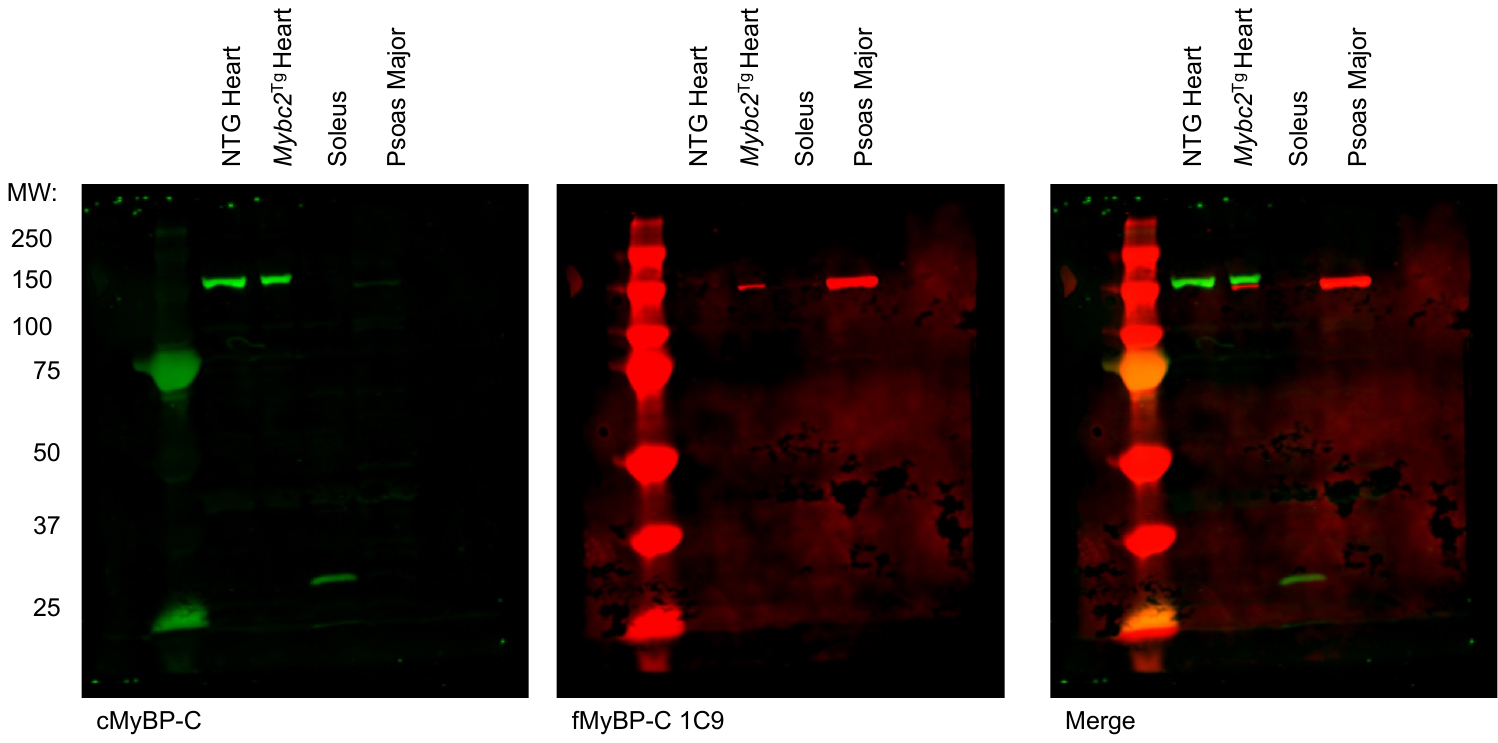

**Supplementary Figure 3. Validation of custom 1C9 monoclonal fMyBP-C antibody.**

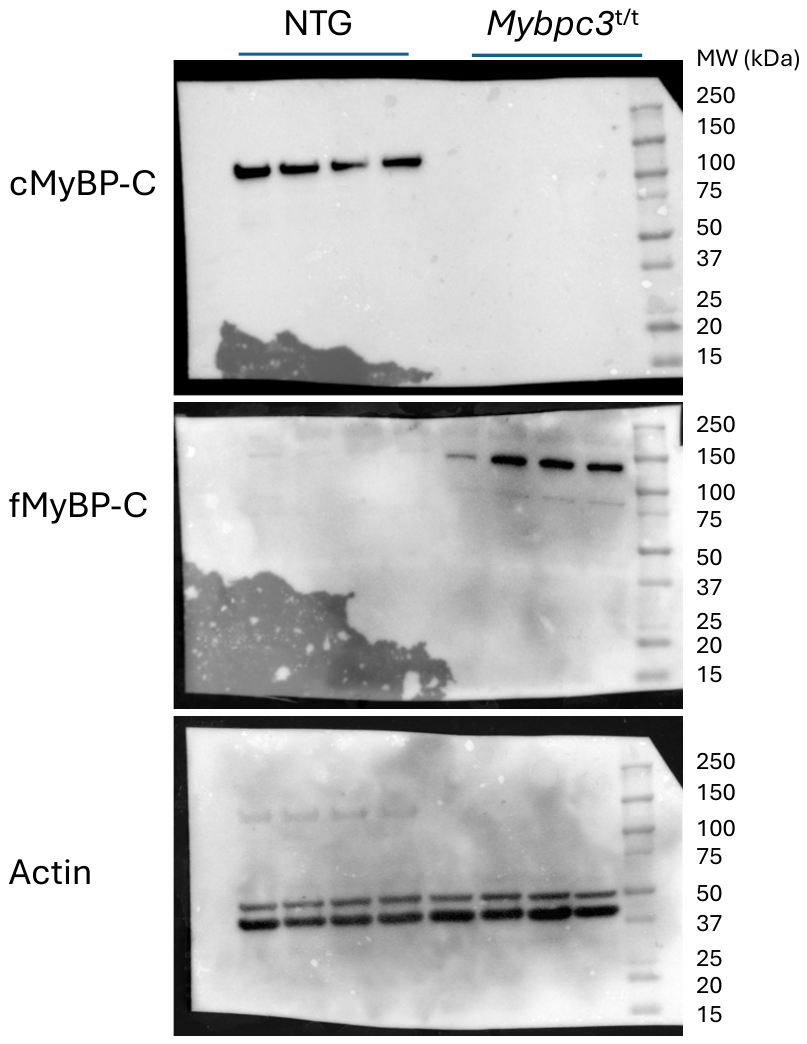

**Supplementary Figure 4. Original Western Blots Corresponding to Figure 1B**

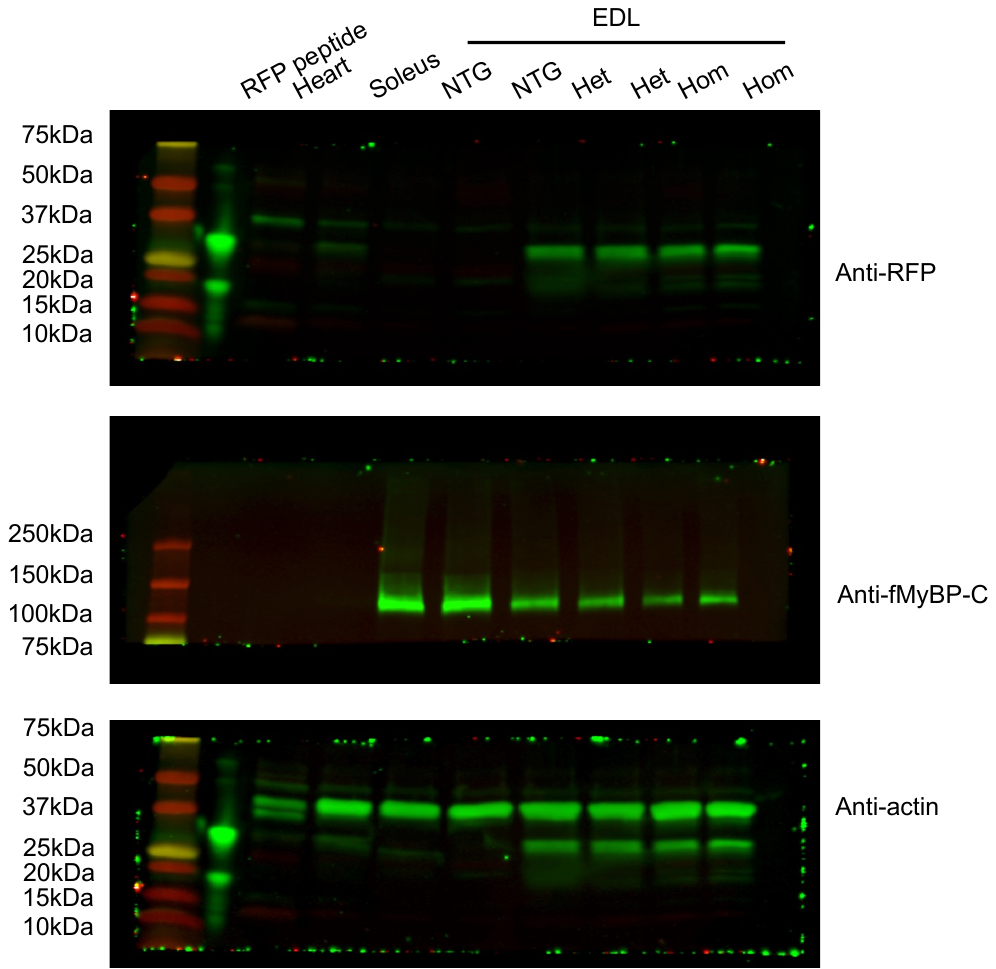

**Supplementary Figure 5. Original Western Blots Corresponding to Figure 1H**

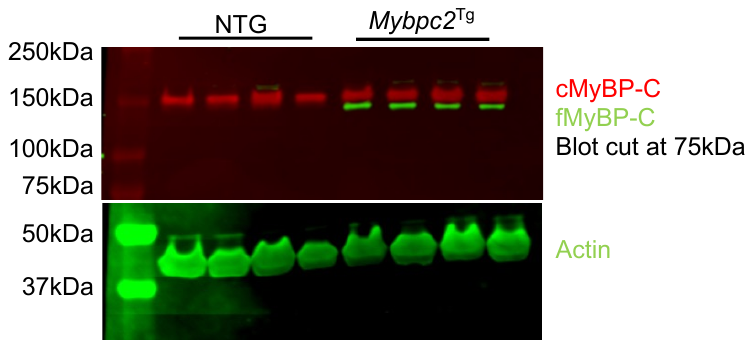

**Supplementary Figure 6. Original Western Blots Corresponding to Figure 2B**

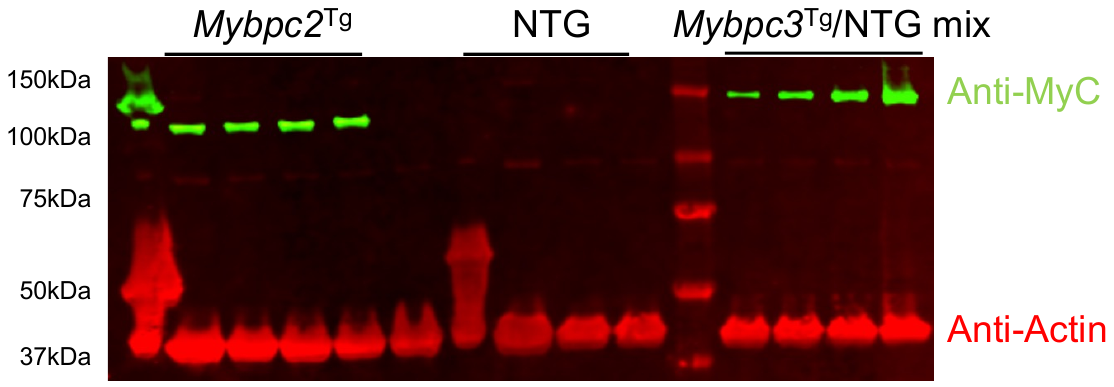

**Supplementary Figure 7. Original Western Blots Corresponding to Figure 2D**

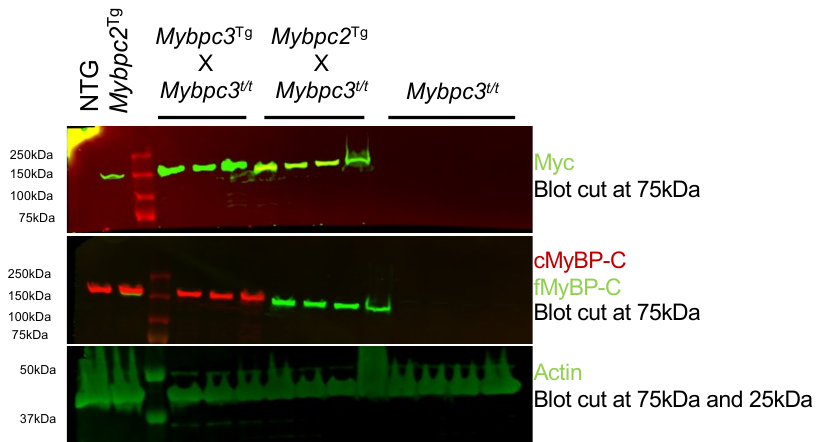

**Supplementary Figure 8. Original Western Blots Corresponding to Figure 4A**

**Supplementary Table 1. Echocardiographic parameters of *Mybpc2*^Tg^ and non-transgenic littermates**

|  | NTG (n=5) | | *Mybpc2*^Tg^ (n=6) | | p-value |
| --- | --- | --- | --- | --- | --- |
|  | Mean | S.E.M | Mean | S.E.M |  |
| HR | 469.40 | 17.90 | 459.50 | 13.49 | 0.6631 |
| LVID;s (mm) | 2.71 | 0.09 | 2.97 | 0.09 | 0.0653 |
| LVID;d (mm) | 3.93 | 0.06 | 4.29 | 0.06 | 0.0026 |
| LV Vol;s (mm) | 27.48 | 2.31 | 34.50 | 2.44 | 0.0697 |
| LV Vol;d (mm) | 67.31 | 2.45 | 83.23 | 2.68 | 0.0020 |
| SV (μl) | 39.85 | 1.51 | 48.70 | 1.80 | 0.0052 |
| EF (%) | 59.36 | 2.31 | 58.65 | 2.13 | 0.8267 |
| FS (%) | 31.16 | 1.55 | 30.91 | 1.52 | 0.9113 |
| CO (ml/min) | 18.79 | 1.16 | 22.32 | 0.77 | 0.0280 |
| IVS;d (mm) | 0.86 | 0.04 | 0.87 | 0.04 | 0.8596 |
| IVS;s (mm) | 1.23 | 0.05 | 1.14 | 0.08 | 0.4219 |
| LVPW;d (mm) | 0.98 | 0.06 | 0.96 | 0.04 | 0.7817 |
| LVPW;s (mm) | 1.20 | 0.06 | 1.23 | 0.06 | 0.7115 |

HR, Heart Rate; LVID, left ventricular internal diameter; LV Vol, left ventricular volume; SV, stroke volume; EF, ejection fraction; FS, fractional shortening; CO, cardiac output; IVS, interventricular septal thickness; LVPW, left ventricular posterior wall thickness; s, systolic measurement; d, diastolic measurement. p-values were calculated by unpaired students t-test.

**Supplementary Table 2. Echocardiographic parameters of Mybpc2^Tg^ (t/t) and *NTG (t/t)* littermates**

|  | Mybpc2^Tg^ (t/t) (n=8) | | *NTG (t/t)* (n=9) | | p-value |
| --- | --- | --- | --- | --- | --- |
|  | Mean | S.E.M | Mean | S.E.M |  |
| HR | 399.84 | 17.85 | 446.44 | 20.56 | 0.086 |
| LVID;s (mm) | 4.56 | 0.17 | 4.02 | 0.10 | 0.021 |
| LVID;d (mm) | 5.26 | 0.15 | 4.80 | 0.09 | 0.030 |
| LV Vol;s (mm) | 96.60 | 8.92 | 71.88 | 4.93 | 0.026 |
| LV Vol;d (mm) | 134.02 | 9.22 | 108.37 | 5.16 | 0.035 |
| SV (μl) | 37.41 | 1.92 | 36.48 | 2.20 | 0.769 |
| EF (%) | 28.52 | 1.97 | 34.09 | 1.85 | 0.045 |
| FS (%) | 13.46 | 1.01 | 16.21 | 0.97 | 0.057 |
| CO (ml/min) | 15.07 | 1.22 | 16.22 | 1.41 | 0.481 |
| IVS;d (mm) | 0.91 | 0.06 | 1.12 | 0.07 | 0.011 |
| IVS;s (mm) | 1.01 | 0.07 | 1.29 | 0.08 | 0.005 |
| LVPW;d (mm) | 0.81 | 0.10 | 1.12 | 0.10 | 0.020 |
| LVPW;s (mm) | 0.91 | 0.11 | 1.29 | 0.13 | 0.011 |
| Corr LVmass (mg) | 157.55 | 10.89 | 199.17 | 12.49 | 0.032 |

HR, Heart Rate; LVID, left ventricular internal diameter; LV Vol, left ventricular volume; SV, stroke volume; EF, ejection fraction; FS, fractional shortening; CO, cardiac output; IVS, interventricular septal thickness; LVPW, left ventricular posterior wall thickness; Corr LVmass, corrected left ventricular mass; s, systolic measurement; d, diastolic measurement. p-values were calculated by unpaired students t-test.
