## Supplementary material for "Fast skeletal myosin binding protein-C expression exacerbates dysfunction in heart failure": Online Methods

***Animal Models***

All experiments were conducted under institutional guidelines and approved by the University of Cincinnati Animal Care and Use Committee for Dr. Sadayappan (AM17-22-11-21-01). To generate cardiac specific over expression of *Mybpc2*, the *Mybpc2* transcript (NM_146189.3) to include an in frame Myc tag replacing the initiating methionine and flanking SalI restriction endonuclease restriction sites at each end. The resulting PCR product was cloned into the pBS-Sk2 + aMHC promoter vector (a gift from Dr. Jeffrey Robbins). After NotI backbone liberation of the vector backbone, the aMHC-*Mybpc2* transgene was injected into FvB/NJ zygotes. Founders and their offspring were screened with the following primers intron spanning primers: 5’-CTGTCAAGGATCTCCCAACTGGA-3’ and 5’-GATATAAGTCTGACGAAGATGGCGGG-3’. *Mybpc3^t/t^* mice have been previously described (1, 2). *Mybpc2*^KO^ mice were generated using CRISPR/Cas9 to induce a deletion between exons 6 and 7 of *Mybpc2*. Two synthetic gRNAs were used: 5’- GAAGCTCTCGAGGCTCTGAC-3’ and 5’- GCAGGCGAGCTGGATTTCAG-3’. Founders were screen with primers 5’-GGAGGAAAGTGAATTGTGAATGC -3’ and 5’-GAGGTGTTGGCACCTTGGCTAC-3’. A founder with a 418bp deletion, corresponding to the production of a 157 amino acid protein product. *Mybpc2^mScarlet^* were also generated using CRISPR/Cas9 to introduce an IRES-mScarlet sequence into the 3’ UTR of *Mybpc2* using guide RNA sequence 5’-CTAAATAAGTCATCAGAACC-3’.

***Transthoracic Aortic Constriction***

Pressure overload of the LV was induced by TAC surgery, as previously described (3). Briefly, animals were anesthetized using isoflurane, intubated, and ventilated. Following sternotomy, a 7-0 silk suture was used to tie down the aorta (between the innominate and left carotid arteries) around a 27-gauge needle (which was subsequently removed) to produce a constriction of uniform and reproducible diameter. Sham-operated animals were subjected to the identical surgical procedures, besides the fact that the suture was not tied to produce an occlusion.

***RNA Isolation***

RNA was isolated as previously described (1, 4). Briefly, a small piece of tissue was homogenized in PureZOL reagent (BioRad). After incubation at room temperature for 5 minutes, phase separation was performed using chloroform. After centrifugation, the aqueous layer was precipitated with ice cold 70% (vol/vol) ethanol. Purification was then performed with using the Qiagen RNeasy mini kit (74104) following manufacturers procedures. RNA concentration and quality was measured using a Nanodrop and electrophoresis using a 3% agarose gel treated with 2% bleach.

***Quantitative Polymerase Chain Reaction***

cDNA was synthesized from 500ng RNA using the iScript cDNA synthesis kit (BioRad, 1708890). Following synthesis, cDNA was diluted 1:10 in nuclease free water and frozen at -20°C. Quantitative PCR was performed using [iTaq Universal SYBR Green Supermix](https://www.bio-rad.com/en-us/product/itaq-universal-sybr-green-supermix?ID=M87FTF8UU) (BioRad 1725121) using the following primers.

*Mybpc2*:5’-CAGTGTGAGGCTGAACTCATT-3’ 5’-GTCACCTTCTCATCAGACACTTC-3’ *Mybpc3*: 5’-GACACCTTCATCTTCCTTCTCTG-3’ 5’-TGAGGAGTGGGTGTTTGATAAG-3’

*Nppa*: 5’-CTGGGACCCCTCCGATAGAT-3’ 5’-TTCGGTACCGGAAGCTGTT-3’

*Nppb*: 5’-AAGGACCAAGGCCTCACAAA-3’ 5’-ACAACAACTTCAGTGCGTTACA-3’

*Actb*: 5’-GGCTGTATTCCCCTCCATCG-3’ 5’-CCAGTTGGTAACAATGCCATGT-3’

*Myh7*: 5’-GGAGAATCAGTCCATCCTCATC-3’ 5’-CCCAATGGCGGCAATAAC-3’

***Protein Isolation***

Myofilament from left ventricular tissue were enriched as described(4). Briefly, left ventricular tissue was minced and homogenized F60 buffer (60mM KCl, 30mM imidazole, 2mM MgCl_2_) containing protease inhibitor, after centrifugation the myofilament-containing pellet was treated with F60 buffer + 1% (vol/vol) Triton X-100 followed by centrifugation three times. Pellets were washed one time with F60 buffer before solubilization in protein solubilization buffer (BioRad 1632145). For total left ventricular homogenates, tissue was homogenized directly in protein solubilization buffer. Protein concentrations were measured using Bradford dye (Thermo Scientific 23200) using bovine serum albumin as a standard.

***Western Blotting***

Proteins were separated by SDS-PAGE using 4-15% Criterion tris-HCl acrylamide gels (BioRad). After electrophoresis, proteins were transferred onto nitrocellulose membranes by wet tank transfer at 300mA for 3 hours with ice cooling. Blots were blocked with 5% skim milk in TBS-T and incubated overnight in primary antibodies at 4°C with rocking. The following primary antibodies used as follows: cMyBP-C (Rabbit polyclonal custom antibody against amino acids 2-14), fMyBP-C (custom mouse monoclonal raised against C1C2 region of mouse fMyBP-C, and Sigma Aldrich SAB2108180), RFP (rabbit polyclonal Rockland, 600-401-379), cMyc (rabbit polyclonal sigma C3956). Following 3 washes with TBS-T, blots were incubated in either Licor IRDye 800CW and 680RD secondary antibodies (1:10,000) or traditional HRP-conjugated secondary antibody for imaging on a Licor Odyssey DLx or BioRad Chemidoc, respectively. Band intensity was quantified using either Licor Image Studio or BioRad ImageLab.

***Cardiomyocyte Isolation and contractility***

Cardiomyocytes were isolated from mouse left ventricles using the Langendorff procedure as described (5). Mice were injected with 0.15ml heparin (1000U/ml) and deeply anesthetized with euthasol (200mg/kg). After excision, hearts were mounted by the aorta on the Langendorff system for retrograde perfusion with a perfusion buffer consisting of NaCl (120 mm), KCl (5.4 mm), MgSO_4_ (1.4 mm), NaH_2_PO_4_ (12 mm), NaHCO_3_ (20 mm), 2,3-butadiene monoxime (10 mm), taurine (5 mm), and glucose (5.6 mm) at 37 °C for 3 min. After this the hearts were perfused for 8-15 minutes with the same buffer but with 0.25mg Liberase (Roche Applied Science) and 12.5 μM CaCl_2_ until the heart became flaccid. The ventricles were removed and dissociated by trituration, filtered, and allowed to pellet by gravity. These cardiomyocytes were resuspended in perfusion buffer with 0.5% BSA followed by sequential restoration of extracellular calcium to a concentration of 25 μM, 100 μM, 200 μM, and finally 1 mm calcium solution. This cell suspension was used for contractility or immunofluorescence. For calcium kinetics, the cells were resuspended in 1.8 mm Ca^2+^-Tyrode's solution containing NaCl (140 mm), KCl (4 mm), MgCl_2_ (1 mm), glucose (10 mm), and HEPES (5 mm), pH 7.4, at room temperature. *Mybpc2*^mScarlet^ cardiomyocytes were immediately observed under the fluorescent microscope to capture the endogenous mScarlet signal. Additionally, the remaining CMs were fixed and used for the immunofluorescent analysis.

***Immunofluorescence***

Cardiomyocytes and hearts were fixed in 4% paraformaldehyde. Hearts were then cryoprotected with sucrose and embedded in OCT. 10μm sections were cut using a cryostat. For immunofluorescent staining, tissue was first permeabilized with PBS containing 0.1% triton X-100, followed by blocking in 5% BSA in PBS containing 0.1% tween-20. Primary antibodies were incubated overnight at 4°C followed by washing and secondary incubation at room temperature for one hour. The following primary antibodies were used: cMyc (rabbit polyclonal sigma C3956), fMyBP-C (rabbit polyclonal Sigma Aldrich, SAB2108180), α-Actinin (mouse monoclonal, Sigma A7811), RFP (rabbit polyclonal Rockland, 600-401-379), cardiac troponin-T (mouse monoclonal, Fisher Scientific MS295P1). DAPI was using for nuclei visualization at a concentration of 1ug/ml. Cell membranes were visualized by staining with Wheat Germ Agglutinin labelled with Alexa Fluor488 (ThermoFisher).

***Mass Spectrometry of tissue proteomics***

Ventricular tissues were snap frozen in liquid nitrogen and stored until analysis. To increase proteome coverage, cardiac proteins were fractionated into cytoplasmic-, myofilament- and insoluble-enriched protein extracts as outlined (6). 100μg of each fraction, quantified by BCA assay was reduced with 1mM TCEP and washed using S-trap mini spin columns (ProtiFi, LongIsland, NY). Each fraction was then subjected to in-column trypsin digestion and eluted. Tryptic peptide concentration was determined using the Pierce Quantitative Colorimetric Peptide Assay Kit (Thermo Fisher Scientific). Samples were run in data independent acquisition mode (DIA) on the Orbitrap Fusion Lumos mass spectrometer equipped with an Easy Spray ion source and connected to Ultimate3000nanoLC system (Thermo Scientific) as described (7). Raw proteomic data files were converted to mzml format using MSConvert. Files were searched using DIANN with label-free quantitation and match-between-runs approach against the complete mouse protein Uniprot database. Identified proteins based on peptides comprised of amino acid sequence unique to a protein, isoform identification was based on an isoform-specific peptide. The mass error tolerances set at 15 ppm for both fragment and intact masses. The library-free identifications are set at <1% false discovery rate (FDR).

The mass spectrometry proteomics data have been deposited to the ProteomeXchange Consortium via the PRIDE [1] partner repository with the dataset identifier PXD047728.

***Cardiac Function Measurement***

Cardiac function was assessed non-invasively by echocardiography using the Vevo 2100 and the 30mHz MS-400 (FUJIFILM VisualSonics Inc., Toronto, Canada) probe as described (1). Briefly, mice were anesthetized with 1-2% isoflurane and chest hair removed. Parasternal long- and short-axis B-Mode and M-Mode images were collected. Images were analyzed using Vevo Lab (FUJIFILM VisualSonics Inc., Toronto, Canada).

Cardiac function was assessed by invasive hemodynamics as previously described (8, 9). Mice were anesthetized with ketamine-thiobutabarbital, and a tracheotomy performed using polyethylene-90 tubing. The right femoral artery and vein were cannulated with polyethylene tubing to measure systemic arterial pressure and delivery drug. To assess myocardial performance, closed chest animals were studied with a high fidelity 1.2F Scisense micromanometer tipped catheter placed retrograde across the aortic valve into the LV via the right carotid artery for pressure. Cardiovascular response to increasing doses of dobutamine were determined during 3-minute constant infusions (0.1 µl/min/gm body weight), and average values determined during the final 30 seconds of each infusion. Online heart rate, telemetry, LV developed pressure, +dP/dt, and -dP/dt were measured using PowerLab (ADInstruments).

***Skinned Fiber Mechanics***

The mechanical properties of chemically skinned fibers were assessed using the 1400A permeabilized fiber system (Aurora Scientific, Aurora, ON, Canada) using protocols identical to those described in (10-12). Frozen left ventricular muscle was thawed and homogenized in a pCa 9.0 relaxing buffer (in mM: 7 EGTA, 100 BES, 0.017 CaCl2, 5.491 MgCl2. 5 DTT, 15 creatine phosphate, and 4.655 ATP; pH adjusted to 7.0 with KOH and ionic strength set to 180 with K-propionate. Triton X-100 was added to a concentration of 1% (vol/vol) to allow skinning overnight on ice. After washing with fresh relaxing buffer, muscle fibers were attached at each end with aluminum t-clips and mounted between the force transducer and length controller of the 1400A system. All measurements were taken at a sarcomere length of 2.1μm and 22°C.

***Skinned Fiber mantATP Turnover Assays***

Skinned fiber mantATP turnover assays were performed as described (10). Left ventricle was skinned on ice for six hours in a buffer consisting of (in mM) 100 NaCl, 8 MgCl2, 5 EGTA, 5 K2HPO4, 5 KH2PO4, 3 NaN3, 5 ATP, 1 DTT, 20 BDM and 0.1% (v/v) Triton X-100 at pH 7, followed by overnight glycerination in a buffer consisting of (in mM): 120 K acetate, 5 Mg acetate, 5 EGTA, 2.5 K2HPO4, 2.5 KH2PO4, 50 MOPS, 5 ATP, 20 BDM, 2 DTT, and 50% (v/v) glycerol at pH 6.8, and after ~17 hours, new glycerinating solution was added. Tissue was stored at -20°C for use within 5 days. Small bundles were dissected and mounted in a flow cell and washed with rigor buffer consisting of (in mM) 120 K acetate, 5 Mg acetate, 5 EGTA, 2.5 K2HPO4, 2.5 KH2PO4, 50 MOPS, and 2 fresh DTT at pH 6.8. Next, rigor buffer containing 250µM mantATP (Sapphire Bioscience, Cat # NU-202L), a fluorescent analogue of ATP, was loaded into the cell chamber. After 60 seconds of imaging, the mant-ATP was replaced with a relaxing solution identical to the rigor solution but containing 4mM ATP. Imaging was carried out for up to 6 minutes. All SRX assays were performed at room temperature (approximately 22°C).

***Development of anti-fMyBP-C monoclonal antibody***

A short-term immunization procedure was applied to immunize an 8-week-old female Balb/c mouse. Recombinant C1C2 domain of fMyBP-C (a.a. 1-337) was expressed from cDNA cloned in pET28a+ plasmid with an N-terminal His-tag in transformed bacterial culture and purified using metal affinity chromatography. 50 μg of the purified C1C2 protein in 100 μL phosphate buffered saline (PBS) was mixed with an equal volume of Freund’s complete adjuvant and injected intraperitoneally and intramuscularly. Ten days after the primary immunization, the mouse was intraperitoneally boosted daily with 100 μg of the C1C2 antigen in 200 μL PBS without adjuvant on two consecutive days. Two days after the final boost, spleen cells were harvested from the immunized mouse for fusion with Sp2mIL-6 mouse myeloma cells (ATCC CRL-2016) using 50% polyethaglycol_3500_ (Invitrogen) containing 7.5% dimethyl sulfoxide as described previously . Hybridomas growing in 96 well culture plates were selected with HAT (0.1 mM hypoxanthine, 0.4 μM aminopterin, 16 μM thymidine) media containing 20% fetal bovine serum and screened using indirect enzyme-linked immunosorbent assay (ELISA) against plate-immobilized C1C2 protein using horseradish peroxidase (HRP)-labeled goat anti-mouse total immunoglobulin second antibody (Santa Cruz). Positive anti-fMyBP-C C1C2 antibody-secreting hybridoma clones were subcloned three times using the limiting dilution method (13) to establish stable cell lines. Culture supernatants of the hybridoma cells was collected and stored in aliquots after lyophilization.

Monoclonal antibody produced by one of the hybridoma lines (1C9) was used in the present study after validation of its specific recognition of fMyBP-C in Western blot of total protein extracts of skeletal and cardiac muscles (Fig. S3).

***Statistical Analysis***

All values are represented as mean ± SEM. statistical analysis was performed using GraphPad Prism. Statistically significant differences between two groups were tested by an unpaired Student’s t-test. One-way ANOVA was used when comparing more than two groups, followed by post hoc analysis with Tukey’s multiple comparison test. All. Statistical significance was defined as p < 0.05.

1.	D. Y. Barefield *et al.*, Ablation of the calpain-targeted site in cardiac myosin binding protein-C is cardioprotective during ischemia-reperfusion injury. *J Mol Cell Cardiol* **129**, 236-246 (2019).

2.	B. K. McConnell *et al.*, Dilated cardiomyopathy in homozygous myosin-binding protein-C mutant mice. *J Clin Invest* **104**, 1235-1244 (1999).

3.	L. C. Green *et al.*, Human antigen R as a therapeutic target in pathological cardiac hypertrophy. *JCI Insight* **4** (2019).

4.	T. L. t. Lynch *et al.*, Amino terminus of cardiac myosin binding protein-C regulates cardiac contractility. *J Mol Cell Cardiol* **156**, 33-44 (2021).

5.	M. Kumar, K. Haghighi, E. G. Kranias, S. Sadayappan, Phosphorylation of cardiac myosin-binding protein-C contributes to calcium homeostasis. *J Biol Chem* **295**, 11275-11291 (2020).

6.	L. A. Kane, I. Neverova, J. E. Van Eyk, "Subfractionation of Heart Tissue" in Cardiovascular Proteomics: Methods and Protocols*,* F. Vivanco, Ed. (Humana Press, Totowa, NJ, 2007), 10.1385/1-59745-214-9:87, pp. 87-90.

7.	L. Ai *et al.*, High-Field Asymmetric Waveform Ion Mobility Spectrometry: Practical Alternative for Cardiac Proteome Sample Processing. *J Proteome Res* **22**, 2124-2130 (2023).

8.	J. Rubinstein *et al.*, Novel role of transient receptor potential vanilloid 2 in the regulation of cardiac performance. *Am J Physiol Heart Circ Physiol* **306**, H574-584 (2014).

9.	S. Sadayappan *et al.*, Cardiac myosin-binding protein-C phosphorylation and cardiac function. *Circ Res* **97**, 1156-1163 (2005).

10.	J. W. McNamara, R. R. Singh, S. Sadayappan, Cardiac myosin binding protein-C phosphorylation regulates the super-relaxed state of myosin. *Proc Natl Acad Sci U S A* **116**, 11731-11736 (2019).

11.	T. Song *et al.*, Fast skeletal myosin-binding protein-C regulates fast skeletal muscle contraction. *Proc Natl Acad Sci U S A* **118** (2021).

12.	R. R. Singh, J. W. McNamara, S. Sadayappan, Mutations in myosin S2 alter cardiac myosin-binding protein-C interaction in hypertrophic cardiomyopathy in a phosphorylation-dependent manner. *J Biol Chem* **297**, 100836 (2021).

13.	J. P. Jin, M. L. Malik, J. J. Lin, Monoclonal antibodies against cardiac myosin heavy chain. *Hybridoma* **9**, 597-608 (1990).
